# Structure-Function Coupling in Highly Sampled Individual Brains

**DOI:** 10.1101/2023.10.04.560909

**Authors:** Aishwarya Rajesh, Nicole A. Seider, Dillan J. Newbold, Babatunde Adeyemo, Scott Marek, Deanna J. Greene, Abraham Z Snyder, Joshua S. Shimony, Timothy O Laumann, Nico U.F. Dosenbach, Evan M. Gordon

## Abstract

Structural connections (SC) between distant regions of the brain support synchronized function known as functional connectivity (FC) and give rise to the large-scale brain networks that enable cognition and behavior. Understanding how SC enables FC is important to understand how injuries to structural connections may alter brain function and cognition. Previous work evaluating whole-brain SC-FC relationships showed that SC explained FC well in unimodal visual and motor areas, but only weakly in association areas, suggesting a unimodal-heteromodal gradient organization of SC-FC coupling. However, this work was conducted in group-averaged SC/FC data. Thus, it could not account for inter-individual variability in the locations of cortical areas and white matter tracts. We evaluated the correspondence of SC and FC within three highly sampled healthy participants. For each participant, we collected 78 minutes of diffusion-weighted MRI for SC and 360 minutes of resting state fMRI for FC. We found that FC was best explained by SC in visual and motor systems, as well as in anterior and posterior cingulate regions. A unimodal-to-heteromodal gradient could not fully explain SC-FC coupling. We conclude that the SC-FC coupling of the anterior-posterior cingulate circuit is more similar to unimodal areas than to heteromodal areas.

**SIGNIFICANCE STATEMENT:** Structural connections between distant regions of the human brain support networked function that enables cognition and behavior. Improving our understanding of how structure enables function could allow better insight into how brain disconnection injuries impair brain function.

Previous work using neuroimaging suggested that structure-function relationships vary systematically across the brain, with structure better explaining function in basic visual/motor areas than in higher-order areas. However, this work was conducted in group-averaged data, which may obscure details of individual-specific structure-function relationships.

Using individual-specific densely sampled neuroimaging data, we found that in addition to visual/motor regions, structure strongly predicts function in specific circuits of the higher-order cingulate gyrus. The cingulate’s structure-function relationship suggests that its organization may be unique among higher-order cortical regions.

## Introduction

The human brain is organized into large-scale brain networks consisting of distributed and functionally connected brain regions which enable perception and cognition (Petersen & Sporns 2015). Distributed brain regions are physically connected via long-range axonal projections grouped into large white matter tracts; such physical connections can be measured in vivo using MRI diffusion tractography techniques and are referred to as structural connectivity (SC; Jeurissen et al., 2019). Additionally, the patterns of associations between these distributed brain regions can be measured as correlations in the spontaneous activity observed in resting-state functional MRI and are referred to as functional connectivity (FC; Stephen, Friston & Squire, 2009).

Knowing how SC supports FC–that is, understanding the nature of structure-function (S-F) coupling—is critical to understanding how the brain’s physical structure supports the integration of sensory representations and execution of cognitive functions, which in turn inform our overall experience (Wang et al., 2018). A deeper understanding of the interaction between SC and FC in healthy normative individuals would subsequently allow for better characterization of dysfunction associated with structural brain injury, such as disconnection syndromes, idiopathic generalized epilepsy, and traumatic brain injury (TBI; Cocchi et al., 2014; Chiang et al., 2015; Kuceyeski et al.,2016, 2019).

Early observations in restricted sets of brain regions favored the idea that functional connectivity relies on the presence of direct, underlying white-matter connections (*for a review, see* Damoiseaux & Greicius, 2009). However, broader testing revealed that this correspondence is not generalizable. At the macroscale, highly integrated functional networks, such as the default-mode network, show weak correspondence with related structural networks (Sarwar et al., 2021). Further, absent, or compromised SC due to brain injury does not necessarily cause reduced FC (O’Reilly et al., 2013; Hawallek et al., 2011; Owen et al., 2013; Tyszka et al., 2011).

Accumulating evidence converges on the observation that FC is not solely dependent on the presence of directly connected white matter tracts. Instead, FC between any two regions is likely to be driven both by tracts directly connecting the regions, as well as through multi-tract (polysynaptic) connections that include other regions (Goñi et al., 2014). Based on this conceptualization, a recent study (Vázquez-Rodríguez et al., 2019) used a multiple linear regression framework to determine how spatial proximity and tract-based SC topology measures — specifically, path length and communicability — predict FC across different brain regions. This model showed high S-F coupling in unimodal brain areas such as visual and motor cortical regions. In contrast, low S-F coupling was found in functionally diverse, multimodal brain areas. These regions are typically recruited for information integration and higher-order cognitive processing. These findings were consistent with the unimodal-to-multimodal axis of cortical organization, providing support to the theory that S-F coupling evolved systematically, in line with cortical expansion (Margulies et al., 2016).

Such approaches that use both direct and indirect SC to model FC provide a useful framework for understanding S-F coupling. However, this model was tested in short-acquisition, group-averaged SC and FC data. Both FC and SC measures are unreliable unless relatively large quantities of within-individual data are collected (Laumann et al., 2015; Gordon et al., 2017, Seider et al., 2022). Further, details of brain organization vary substantially across individuals (Laumann et al., 2015; Gordon et al., 2017, Braga & Buckner, 2017, Somers et al., 2021). Across different participants, a given atlas coordinate cannot be expected to be a part of the same brain network, with the same patterns of FC and SC. Averaging across this variation obscures details of brain organization that are spatially variable across individuals, preventing accurate evaluation of S-F coupling. Indeed, the primary sensory/motor regions which have previously exhibited the highest S-F coupling (Vázquez-Rodríguez et al., 2019) are those with the lowest inter-individual variation (Gordon et al., 2017).

Here, we adapted the Vázquez-Rodríguez model to a precision neuroimaging approach — employing extended-acquisition, within-individual, surface-mapped data to reduce variation due to noise, inter-individual variability, and uncertainty about gray-white boundaries.

We collected 78 minutes of diffusion-weighted MRI for SC and 360 minutes of resting state fMRI for FC from three individuals (27–35 years). We optimized computation of FC by using a surface-based approach, which enables more precise spatial localization of cortical BOLD signals than traditional volume-based approaches (Coalson, Van Essen, & Glasser, 2018). We optimized computation of SC by using a probabilistic tractography (versus a deterministic) approach (Pai, Muzik, & Hua, 2008), coupled with an optimized method to accurately estimate white-matter fibers (Seider et al., 2022). Using this model, we characterized S-F coupling across the individual brain tested if S-F coupling can explain affiliation with individual-specific resting-state networks. We also evaluated S-F coupling patterns for each SC measure. Collectively, our study maximizes our ability to observe S-F coupling at the individual level, and thus represents a critical step toward establishing precision individual patterns in S-F coupling.

## Results

### In individual brains, SC does not consistently vary with FC

For some regions of the individual brain, functional and structural connectivity patterns were well matched, with structural connections linking functionally connected regions. For example, in Participant 01, a posterior cingulate seed exhibited both functional and structural connectivity to medial parietal cortex, medial frontal cortex, and retrosplenial cortex (Fig 1; high S-F coupling). However, for other regions, structural connectivity diverged substantially from functional connectivity. For example, in the same participant, a lateral prefrontal cortex seed exhibited both functional and structural connectivity to anterior lateral frontal cortex and lateral temporal cortex. However, FC with this seed was stronger with posterior superior frontal sulcus, intraparietal sulcus, and anterior insula. These regions were not linked to the frontal cortex seed by the observed SC, suggesting less robust S-F coupling (Fig 1; low S-F coupling).

**Figure 1:**
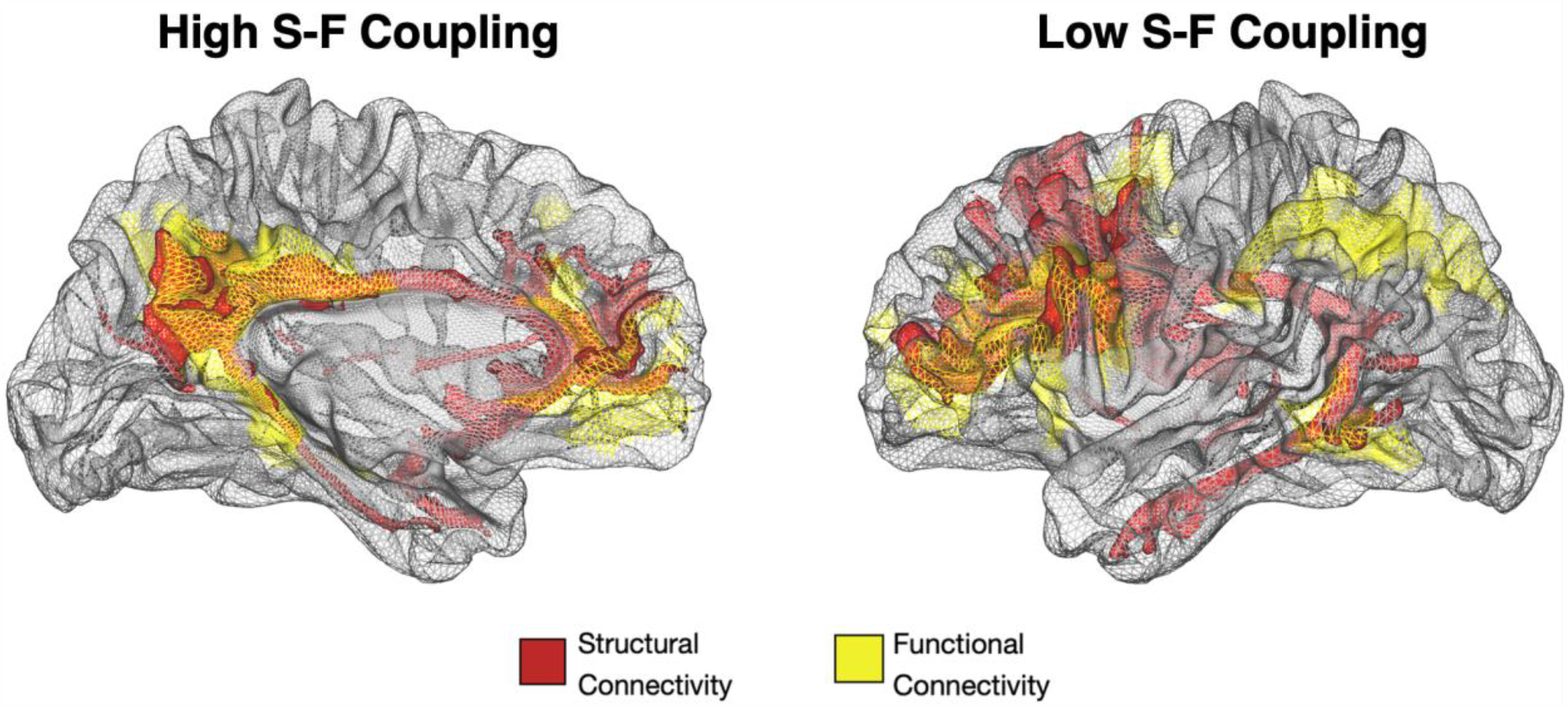
Structural connectivity (SC) and functional connectivity (FC) relationships in a single participant (participant 01). Left: SC and FC were computed from a seed in posterior cingulate cortex. Both FC and SC exhibited connectivity to the posterior cingulate cortex, anterior cingulate and medial frontal cortex, and retrosplenial cortex, suggesting strong S-F coupling. Right: SC and FC were computed from a seed in posterior inferior frontal sulcus. Strong FC was observed with posterior superior frontal sulcus, intraparietal sulcus, and anterior insula. These regions were not linked to the frontal cortex seed by the observed SC, suggesting less robust S-F coupling. For visualization, functional connectivity was thresholded to retain connections with Z(r) > 0.25, while structural connectivity was thresholded to retain tracts with total streamline density > 80,000.

The Vázquez-Rodríguez et al., (2019) multiple linear regression framework well-represented the conceptualization that FC may be driven by spatial proximity (i.e., Euclidean distance) and by multi-tract connections. Such polysynaptic pathways could include the shortest and densest one-way tract connections (i.e., path length) and pathways that include indirect connections with opportunity for back-and-forth (i.e., communicability). We tested this model in each individual-specific parcel.

Across parcels, the mean total variance in FC patterns explained by SC did not exceed 7% in any participant (Figure 2), compared to the 30% observed at the group-level (Vázquez-Rodríguez et al., 2019). These results suggest that spatial proximity, as well as unidirectional and bidirectional communication pathways, only partially explain FC at an individual level (Figure S2). Similar results were observed with an alternate parcellation approach (Figure S4).

**Figure 2:**
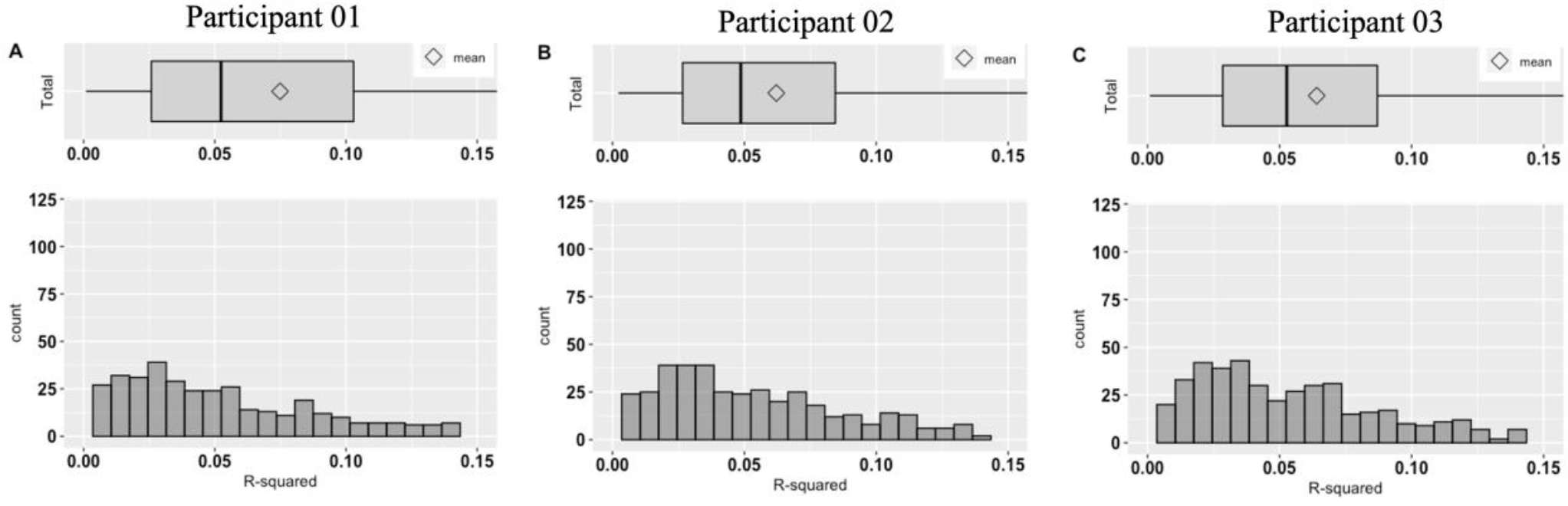
Combined variance contributions by Euclidean distance, path length, and communicability toward FC represented on box and histogram plots. ***Panels A-C***: *Boxplot and histogram plot for total variance contribution by Euclidean distance, path length, and communicability toward FC for participant 01(A)*,*02(B), and 03(C) respectively. Bins on histogram plot represent number of individual-specific parcels in each bin. Across all panels, R*^*2*^ *values were scaled to 0*.*15 % of variance explained by SC toward FC, for maximum comparability across participants*.

### In individual brains, S-F coupling is strongest in unimodal visual and motor areas, as well as in multimodal cingulate areas

At the individual-level, SC explained relatively high variance in FC within specific areas. S-F coupling was strong in unimodal primary visual (20-43%) and motor cortex (21-27%) (Figure 3), consistent with the S-F coupling pattern noted at the group-level in these regions (Vázquez-Rodríguez et al., 2019). Notably, S-F coupling was also relatively strong in heteromodal anterior cingulate (22-29%) and posterior cingulate cortical areas (14-24%) (Figure 3), inconsistent with the S-F coupling pattern noted at the group-level in these regions (Vázquez-Rodríguez et al., 2019). Similar individual-level patterns in S-F coupling were independently observed for all three coupling metrics employed here, including Euclidean distance, path length, and communicability (Figure S3). These results held for parcels obtained using the local gradient approach (Figure S4).

**Figure 3:**
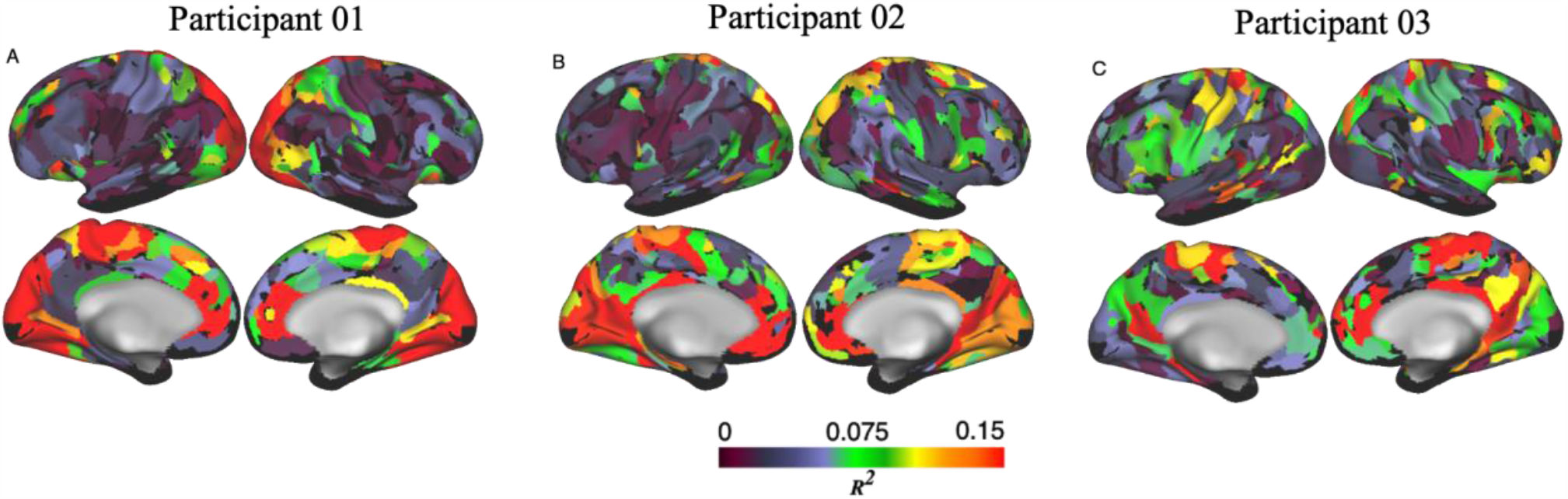
Combined variance contributions by Euclidean distance, path length, and communicability toward FC represented on individual brains. ***Panels A-C: Total*** *variance contribution by Euclidean distance, path length, and communicability toward FC for participant 01(A),02(B), and 03(C)*. **Variance contributions are mapped onto each participant’s specific brain organization, derived using the Infomap community-driven algorithm**. *R*^*2*^ *values were thresholded to 0*.*15 for easy comparison across participants. Black color indicates R*^*2*^ *values = 0*.

At the network level, contributions of SC toward FC were highest for visual and supplementary motor networks, with notable contributions toward the salience network (driven by anterior cingulate cortex) and parietal memory network (driven by posterior cingulate cortex; Figure S5).

Given that observed patterns of S-F coupling across the brain have been framed within theories of unimodal-to-heteromodal cortical expansion (Vázquez-Rodríguez et al., 2019), we tested the association between structure–function R^2^ for a given region and its position along the macroscale unimodal-heteromodal gradient. This association was significant but only weak-to-moderate (Participant 01: *ρ*=-0.33, p<0.001, significant at *α*=0.05; Participant 02: *ρ*=-0.12, p<0.02, significant at *α*=0.05; Participant 03: *ρ*=-0.19, p<0.001, significant at *α*=0.05). Using the map of residuals from this association, we sought to determine which regions violated S-F coupling patterns as would be predicted by the unimodal-heteromodal gradient. We observed mismatches between the S-F coupling pattern and the principal gradient in the anterior and posterior cingulate cortex, where S-F coupling was much stronger than was predicted by the gradient (Figure 4). Taken together, these results suggest that in densely sampled individuals, S-F coupling does not strictly follow the unimodal-multimodal gradient.

**Figure 4:**
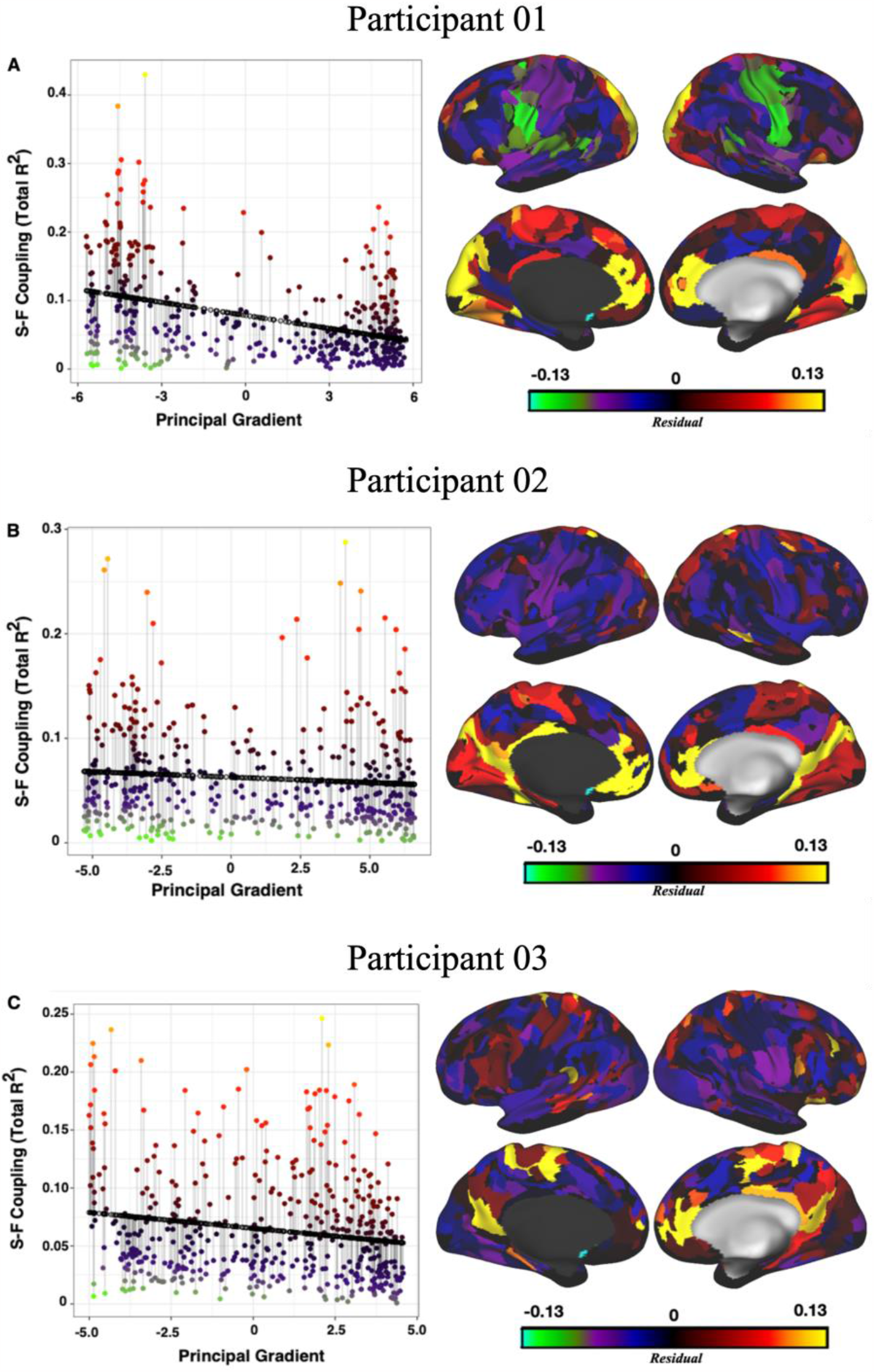
Association between structure–function R^2^ for a given region and its position along the macroscale unimodal-heteromodal gradient. Left: The y-axis represents the strength of S-F-coupling as the total R2 calculated using the multiple linear regression (Vázquez-Rodríguez et al. 2019) approach. The x-axis represents the first principal gradient values obtained using the diffusion embedding (Margulies et al., 2016) procedure. Right: Residuals from the regression plot. PCC and ACC regions exhibited much larger S-F coupling than would be expected given their position on the principal gradient. Residuals are thresholded to values between −0.13 and 0.13, for easy comparison across participants.

## Discussion

We evaluated whether there is a systematic way in which S-F coupling occurs across the brain, in highly sampled individuals. We found that S-F coupling was highest in unimodal visual, motor, and supplementary motor areas and their associated networks. These findings are broadly consistent with prior work using this S-F coupling model (Vázquez-Rodríguez et al., 2019).

However, we also showed that the similarity between distributions of S-F coupling and the individual-specific unimodal-to-heteromodal gradient (Margulies et al., 2016) was relatively weak. This finding is notable, as one major hypothesis for S-F coupling patterns previously observed at the group level has been that S-F coupling evolved systematically, in line with cortical expansion (Vázquez-Rodríguez et al., 2019). Specifically, phylogenetically older regions along the primary sensory/motor areas have dedicated input-output streams and functional responsibilities. Consequently, such regions should show the highest correspondence between structural and functional connectivity. As the cortical mantle expanded further anterior or posterior to these sensory/motor areas, the newly developed regions became untethered from structural constraints and assigned functions. Consequently, such regions are expected to show weaker correspondence between structural and functional connectivity. Here, we show that these expectations do not hold in precision-mapped, individual-specific data, as some association cortex regions exhibit S-F coupling as strong or stronger than primary cortices.

We specifically showed that the divergence with the gradient hypothesis was driven by stronger than expected S-F coupling in multimodal anterior and posterior cingulate cortices (ACC and PCC)—regions which notably exhibit the greatest distance from primary cortices (Margulies et al., 2016). This individual-specific finding is in stark contrast to group-level observations (Vázquez-Rodríguez et al., 2019).

One hypothesis for the strong S-F coupling in the ACC is the presence of Von-Economo neurons (VEN) in this region. It has been suggested that the large size and simple dendritic structure of these projection neurons allow the rapid transfer of information (Allman et al., 2010, Stevens, Hurley, &Taber, 2011). Supporting the possibility that VEN may be involved, we found relatively high S-F coupling in the salience network, a network that maps onto VEN-rich regions (Supplementary Figure 2, Seeley, 2019). Meanwhile, the PCC has been characterized as the ‘structural core’ of the brain (Hagmann et al., 2008). This region shows the highest degree of connectivity in diffusion imaging-based fiber-tracking and has been associated with a wide range of functions (Pearson et al., 2011). One of its most obvious connections is with the ACC via the cingulum bundle. The cingulum bundle may have few crossing fibers and provide a better estimate of SC in the PCC and ACC than in other areas.

Provocatively, these findings suggest that the structure and function of the ACC-PCC circuit may be organized more similarly to primary cortex than to heteromodal association cortical areas. This observation converges with prior work describing functional connectivity-based brain network organization using network visualizations (Power et al., 2011, Gordon et al., 2017, Gordon et al., 2018), which consistently identify both Default and Sensory-Motor regions as being relatively isolated, on the exterior of whole-brain network graphs, while other association cortex regions are more central within the graph. Beyond functional MRI studies, descriptions of the sensorimotor-association axis in anatomical modalities (e.g., intracortical myelin, cortical thickness, allometric scaling, macaque-to-human cortical expansion, and even gene expression patterns) have suggested that PCC in particular may be more similar to sensorimotor cortex than to association cortex (Syndor et al., 2023). The present work supports that concept.

We observed that S-F coupling is weaker at the individual level than has been observed at the group level (Vázquez-Rodríguez et al., 2019). These results are in line with previous work with standard data collections, which show markedly lower agreement between structural and functional connectivity in individuals than in group-level analyses (*see review by* Strathoff et al., 2019). Specifically, in studies that have used comparable approaches as ours (participant-level, probabilistic fiber-tracking, and full functional correlation matrices), the R^2^ values have been much lower (R^2^ range: 0.004-0.006; Horn et al., 2013). Such findings have been theorized to be due to lower signal-to-noise ratios at the individual level (e.g., Skudlarski et al., 2008, Strathoff et al., 2019). However, the precision functional mapping approach used here maximizes reliability of signals in each individual (Gordon et al., 2017, Seider et al., 2022). Thus, lower S-F coupling in individuals is not readily attributable to noise, but should instead be considered close to the true S-F relationships observable in individuals.

We maximized our ability to observe S-F coupling in individuals by employing optimized techniques. First, we used surface-based data (versus the volumetric data used by Vázquez-Rodríguez et al., 2019) to compute FC. Surface-based approaches enable more precise spatial localization of cortical BOLD signals than traditional volume-based approaches (Coalson, Van Essen, & Glasser, 2018). Second, we used a probabilistic (versus a deterministic) tractography approach to compute SC. Probabilistic tractography may facilitate more accurate tracking of white matter fiber courses in complex neighborhoods, such as the branching and crossing fibers that are present in multimodal areas (Pai, Muzik, & Hua, 2008). We enhanced the accuracy of our tractography by employing a recently published crossing-fiber estimation method with higher reliability of crossing-fiber estimation than contemporary algorithms (BaMM, Bayesian Multi-tensor Model-selection; Seider et al., 2022). Third, we investigated our tract-based SC measures — i.e., path length and communicability — using a weighted approach. Weighted measures carry additional information about connection strengths, allowing the accurate representation of SC topology (since the influence of weak links is minimized and the role of strong connections appropriately acknowledged; Rubinov & Sporns, 2010). Finally, we used LMG to estimate unique R^2^ contributions and gauge relative importance of each SC variable toward FC. This statistical method is considered superior to traditionally applied stepwise-regression models (Grömping, U., 2007), which decompose R^2^ to determine relative importance (e.g., Vázquez-Rodríguez et al., 2019). The latter requires a meaningful prior knowledge of the natural order among the SC variables (since R^2^ contributions are obtained by the sequence in which regressors are entered into the regression model; Grömping, U., 2007). The approach adopted by LMG also employs sequential R^2^, but it takes care of the dependence on orderings by averaging over orderings (using simple unweighted averages). Consequently, we were able to generate comparable R^2^ brain maps and evaluate contributions toward networks for each SC variable more accurately.

In this study, we conceptualized the SC-FC relationship as one in which FC depends on SC (Straathof et al., 2019). Accordingly, we acknowledge that the presence of a functional connection may depend on spatial proximity (i.e., Euclidean distance) as well as the presence of a direct (i.e., path length) or indirect structural connection (i.e., communicability). However, our data suggest that the strength of a functional connection does not need to be directly related to the strength of structural connections, as expressed by Euclidean distance, path length, and communicability metrics. It may require other structural connectivity metrics to accurately capture information flow (Straathof et al., 2019). Finally, recent work has suggested that FC may be more strongly influenced by geometric brain shape than by SC (Pang et al., 2023).

In conclusion, we observed that in densely sampled individuals, FC is most strongly explained by SC in visual and motor systems, as well as in anterior and posterior cingulate regions.

## Methods

### Participants

Three healthy, right-handed adult volunteers were recruited as part of a study that used an arm immobilization (“casting”) intervention to induce brain plasticity (Newbold et al., 2020, Seider et al., 2021). The first participant (Participant 01) identified as a 35-year-old male, the second participant (Participant 02) identified as a 25-year-old female, and the third participant (Participant 03) identified as a 27-year-old male.

The data employed here was collected either prior to the immobilization intervention (Participants 02, 03) or two years afterwards (Participant 01). Data from the Cast-Induced Plasticity experiment is publicly shared through the OpenNeuro database (https://openneuro.org/datasets/ds002766/versions/3.0.2). The study was approved by the Institutional Review Board at the Washington University School of Medicine and all participants provided informed consent for all aspects of the study.

### MRI Acquisition

Participants were scanned daily at a consistent hour for 12 sessions prior to the intervention.

#### T1- and T2-weighted structural images

Participant 01’s T1 and T2-weighted structural images were acquired on a Siemens Trio 3T scanner. Across the 12 sessions, a total of four T1-weighted MP-RAGE (sagittal, 224 slices, TE=3.74ms, TR=2400ms, flip angle=8°, 0.8×0.8×0.8mm voxels) and four T2-weighted spin-echo images (sagittal, 224 slices, TE=479ms, TR=3200ms, flip angle=120°, 0.8×0.8×0.8mm voxels) were collected.

For Participants 02 and 03, these images were acquired on a Siemens Prisma 3T scanner. Each session included the collection of a T1-weighted MP-RAGE (sagittal, 208 slices, TE=2.22ms, TR=2400ms, flip angle=8°, 0.8×0.8×0.8mm voxels) and a T2-weighted spin-echo image (sagittal, 208 slices, TE=563ms, TR=3200ms, flip angle=120°, 0.8×0.8×0.8mm voxels).

#### Diffusion-weighted structural images

In each session, a diffusion-weighted sequence was acquired on the Siemens Prisma 3T scanner (single-shot echo planar, axial, 75 slices, TE=83ms, TR=3500ms, 2×2×2 mm voxels, four shells with b-values 250, 500, 1000, and 1500 s/mm^2^, 96 encoding directions). Acquisition of this sequence was only omitted for three sessions in one participant (Participant 01).

#### Resting-state functional images

In each session, we acquired resting-state functional images on the Siemens Prisma 3T scanner. Every session included a 30-minute resting-state fMRI scan collected as a blood oxygen level-dependent (BOLD) contrast sensitive gradient echo-planar sequence (TE=33ms, TR=1100ms, flip angle=84°, 2.6×2.6×2.6mm voxels, multiband 4 acceleration). During this scan, participants were instructed to hold still and look at a white fixation crosshair presented on a black background. A pair of spin echo EPI images were acquired with identical geometrical parameters to the BOLD data but opposite phase encoding directions (AP and PA) to correct spatial distortions.

### Data Processing

#### T1- and T2-weighted structural images

FSL Fast (Zhang et al., 2001) was used to correct gain field inhomogeneity in T1- and T2-weighted images. The 4dfp MRI software package was used to register them to the 711-2B Talairach atlas space (https://readthedocs.org/projects/4dfp/). Mean T1- and T2-weighted images (T1w and T2w) were calculated by co-registration and averaging across acquisitions from the 12 sessions for each participant. The mean T1w image was run through the Freesurfer pipeline (version 5.3, Fischl, 2012) to create anatomical segmentations and 3D surface cortex models.

#### Diffusion-weighted structural images

Acquisition time for each DWI scan was 6.5 min. Across scanning sessions, the total number of DWI scans collected for the three participants were 9, 12, and 14 (i.e., 864, 1152, and 1440 total encoding directions, respectively).

To each DWI, we applied FSL’s Eddy correction and Topup to correct for current-induced and susceptibility-induced distortions (Andersson and Sotiropoulos, 2016; Smith et al., 2004).

FSL’s Eddy correction simultaneously calculates movements on the DWI image, relative to the first volume. Using this information, we removed volumes with framewise displacement > 0.5 mm (Baum et al., 2018). The gradient vectors for each resultant image were concatenated and affine registered to the participant’s mean T1w image, followed by atlas registration. FSL’s tool DTIFIT was used to compute diffusion tensor maps (Jenkinson et al., 2012) and averaged across sessions.

FSL’s Eddy correction also generates rotation corrected b-vectors. These b-vectors were used to estimate tracts using the Bayesian Multi-tensor Model-selection (BaMM) method for fiber estimation (Seider et al., 2022). We have shown that BaMM produces tract angle estimates for tractography that are more precise and reliable than other approaches (Seider et al., 2022).

#### Resting-state functional images

Processing for the resting-state fMRI data has been previously described (Newbold et al., 2020). Briefly, fMRI processing involved correction for odd vs. even slice-intensity differences, across-run intensity normalization, and within- and between-run head motion. The resultant fMRI image was resampled into an isotropic 2-mm atlas space through the following transforms: native space mean resting state image → T2w → T1w → 711-2B template (Jenkinson et al., 2012). The transformation into atlas space took place in a single step and was combined with the registration of session-specific field maps to correct for susceptibility distortion. Subsequent BOLD sessions were linearly aligned to the participant’s first session data.

The 711-2B-transformed image was further preprocessed to reduce variance unlikely to reflect neural activity. Specifically, the 6 rigid-body parameters—obtained from the motion correction method above—were low pass filtered to retain frequencies <0.1 Hz. This filtering process was used to remove respiratory artifacts (Fair et al., 2020). Gray-ordinate image intensity plots (Power, 2017) were visually checked to confirm artifact removal.

Motion-contaminated frames were then flagged, using both outlying values of frame-wise displacement (FD) and variance of derivatives (DVARS) measures (Power et al., 2012), with thresholds set individually for each participant (FD threshold for Participant 01=0.2mm, Participant 02 and 03=0.1mm, DVARS threshold for all participants=6% rms).

Censored frames were replaced by linearly interpolated values. The fMRI time-series was then temporally bandpass filtered (retaining frequencies 0.005-0.1 Hz) and re-censored afterward.

The filtered fMRI image was denoised by regressing out the 6 movement parameters, the global signal averaged over the brain, and orthogonalized waveforms extracted from the ventricles, white matter, and extra-cranial tissues (excluding the eyes).

The denoised image was mapped to Freesurfer-based segmentations and mid-thickness surface models (using the “-ribbon-constrained” method). The different surfaces from these procedures (pial, white matter, and mid-thickness) were brought into register with each other in fs_LR space (Van Essen et al., 2012), and then resampled to the computationally tractable resolution of 32k vertices using Connectome Workbench command line utilities. This surface-mapped data was then combined with data sampled from the individual-specific subcortex (accumbens, amygdala, caudate, hippocampus, pallidum, putamen, and thalamus), and cerebellum to form a Connectivity Informatics Technology Initiative (CIFTI) formatted image. The CIFTI image was spatially smoothed with geodesic (for surface data) and Euclidean (for volumetric data) Gaussian kernels (σ = 2.55 mm; Glasser et al., 2013).

#### Generation of cortical seed regions

Cortical seeds for FC and SC analyses were derived from a graph-theory-based Infomap algorithm for community detection (Rosvall and Bergstrom, 2008, Power et al., 2011) applied to the fMRI data. To this end, we performed the following steps for each participant.

First, we composed the cross-correlation matrix of the surface-mapped fMRI time courses from all brain vertices (on the cortical surfaces) and voxels (in subcortical structures), across sessions.

Second, we estimated the geodesic distance between each vertex and the Euclidean distance between each voxel.

Third, any correlations between vertices/voxels with geodesic/Euclidean distance <= 30 mm was set to zero to avoid basing network membership on spatial smoothing. Inter-hemispheric connections between the cortical surfaces were retained since smoothing was not performed along the mid-sagittal plane.

Fourth, the resultant matrix was thresholded to retain at least the strongest 0.1% of connections to each vertex and voxel (Gordon et al., 2020; 2022). The thresholded matrices were used as inputs for the Infomap algorithm, which calculated community assignments separately for each threshold. The resulting communities represent subnetworks in the brain. The discrete surface-mapped cortical clusters of each subnetwork resulting from this procedure were taken as the cortical regions of interest for FC analyses.

Tractography approaches for SC calculation struggle to accurately track into cortex (Reveley et al., 2015; Van Essen et al., 2013). Thus, tracking streamlines within white matter is preferred to tracking in grey matter (Jackson et al., 2020). Thus, to generate SC seeds for these cortical regions of interest, each cortical seed was projected down into the white matter directly below each cortical seed. Specifically, for each vertex on the fs_LR surface, we computed the 3D vector between corresponding points on the fs_LR_32k pial and the gray-white surfaces, and we extended that vector an additional 2 mm beyond the gray-white surface to create a lower surface, following Gordon et al., 2023. A seed was defined as the region bounded by vertex coordinates of the 2mm-under surface that mirrored the vertex coordinates of the parcel on the fs_LR pial surface above. This resulted in originally cortical seed regions mapped to a lower surface within white matter that is exactly in register to the existing fs_LR_32k surfaces on which the functional data is mapped. Finally, seed regions were projected back into 2mm isotropic volumetric diffusion space and thresholded at values > 0.5 (i.e., thresholded at a > 50% probability that a given voxel is part of that seed region), to minimize spillage of the seed region into other (projected) seeds.

We excluded all subcortical regions from our analysis. Subcortical regions have lower signal-to-noise ratio relative to cortical regions. As a result, they consistently show weaker functional connectivity, which could erroneously influence S-F coupling estimation. The exclusion of subcortical regions is consisted with previous work exploring S-F coupling (Vázquez-Rodríguez et al., 2019).

#### Network identity of seed regions

Finally, we determined the canonical large-scale network identity of each subnetwork using the individual-specific network matching approach described in (Gordon et al., 2017c). Briefly, the Infomap algorithm was applied to each subject’s correlation matrix thresholded at a range of edge densities spanning from 0.01% to 5%. At each threshold, the algorithm returned community identities for each vertex and voxel. Communities were labelled by matching them at each threshold to a set of independent group average networks described in (Gordon et al., 2017c). The matching approach proceeded as follows: 1) At each density threshold, all identified communities were compared to the independent group networks using the Jaccard Index of spatial overlap; 2) The community with the best-match (highest-overlap) to one of the independent networks was assigned that network identity, and then not considered for further comparison with other independent networks within that threshold. Matches lower than Jaccard = 0.1 were not considered (to avoid matching based on only a few vertices). Matches were first made with the large, well-known networks (Default, Lateral Visual, Motor Hand, Motor Mouth, Cingulo-Opercular, Fronto-Parietal, Dorsal Attention), and then to the smaller, less well-investigated networks (Ventral Attention, Salience, Parietal Memory, Contextual Association, Medial Visual, Motor Foot). 3) In each individual and in the average, a “consensus” network assignment was derived by collapsing assignments across thresholds, giving each node the assignment it had at the sparsest possible threshold at which it was successfully assigned to one of the known canonical group networks.

### Analysis

First, we evaluated S-F coupling based on the Vázquez-Rodríguez multiple linear regression model. Second, we evaluated whether S-F coupling follows the principal unimodal-heteromodal gradient as conceptualized by Margulies et al. (2016). To this end, the following variables were computed.

#### Euclidean Distance

Euclidean distance —used to index spatial proximity — was calculated based on the distance between the centers of the cortical seed regions identified above.

#### SC metrics: Weighted Path Length and Weighted Communicability

All variables were estimated using Matlab (version: R2020b). SC matrices were constructed using probabilistic tractography, which quantifies the number of streamlines from a given seed parcel to another target parcel (probtrackx; Behrens et al., 2007). We used the distance correction option to correct for the fact that streamline counts systematically drop with distance from the seed. Further, we note that individual-specific parcels can vary in size, and larger seed parcels cause more tractography streamlines to be initiated. Thus, larger parcels systematically produce higher streamline counts than smaller ones. To correct for this size-dependent tract count, we divided the streamline counts by the total number of streamlines generated for each seed-to-target tracking.

The above procedure resulted in a parcel-to-parcel SC matrix for each participant, where each row represented the proportion of streamlines leaving a seed *i* that reach any of the other targets *j*. Since tractography is dependent on seeding location, the streamline density from *i* to *j* is not necessarily equal to that from *j* to *i*. We therefore defined the undirected connectivity as the average of the streamline densities *ij* and *ji*, which generated an undirected weighted connectivity matrix, SC_weighted_. This SC_weighted_ connectivity matrix was used to create the independent tract-based predictors, i.e., the weighted path length and communicability matrices.

Weighted path length — defined as the minimum number of tracts with maximal densities connecting seed region *i* to seed region *j* — was computed by first converting the SC_weighted_ matrix into a matrix of SC_length_, where 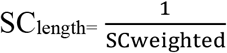. The SC_length_ matrix was used to generate a SC_Path Length_ matrix (using Djikstra’s algorithm as implemented in the Brain Connectivity Toolbox, https://sites.google.com/site/bctnet/?pli=1) such that shortest weighted paths had the minimal weighted distance (but not necessarily the minimal number of edges).

Weighted communicability — defined as the total number of (direct and indirect) tract connections connecting seed region *i* to seed region *j* as well as seed region *j* to *i*— was computed by first acknowledging that the SC_weighted_ matrix encodes direct connections. From this standpoint, the extent to which brain regions that are connected indirectly can be quantified as: 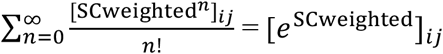. In this context, [SCweighted]^*n*^ characterizes all physical connections between *i* and *j* that traverse *n* tracts. Physical connections that are traversed over *n* tracts are weighted by a factor of 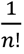, so that longer tract connections contribute less to the sum, while direct connections contribute more. In a weighted matrix, this formulation is modified as:

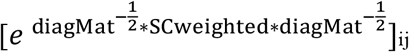, where *diagMat* is the diagonal matrix, and 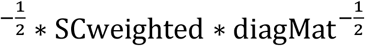 is used to normalize the undue effects of parcellations with high strength (Fornito, Zalesky, & Bullmore, 2016; Worrell et al., 2017).

*FC*

The resting-state FC matrix was computed by generating a set of zero-lag Pearson correlations between time courses of each pair of cortical parcels. These correlations were then fisher-transformed to improve normality.

##### 1. Evaluation of S-F coupling based on the Vázquez-Rodríguez model

To determine the extent to which each parcel-seeded structural metric contributed toward the whole-brain FC pattern of that parcel, we used a multilinear analytic model in R (R studio version: 2021.09.0; R version: 4.1.1). For each individual-specific parcel, the multilinear regression model was constructed as follows:

FC_parcel_ = ***β***_**0**_ + ***β***_**1**_*Euclidean Distance*_*parcel*_ + ***β***_**2**_*Weighted Path Length*_*parcel*_ + ***β***_**2**_*Weighted Communicability*_*parcel*_, where FC_parcel_ is the set of functional connections between a given parcel and every other parcel for a given participant. The independent variables, namely, Euclidean distance, weighted path length, and weighted communicability are calculated for that parcel and every other parcel (Vázquez-Rodríguez et al., 2019).

We optimized this linear regression model by taking the following steps. First, to increase functional specificity, only seed regions separated by a Euclidean distance > 20 mm were considered (i.e., for all variable matrices, if seed-to-seed Euclidean distance ≤ 20 mm, the corresponding cell values were set to 0). Second, we standardized our independent variables (i.e., rescaling by z-scores). Third, we calculated the percent contribution to variation in FC of each independent variable in our model. Traditional approaches to estimating variable contributions toward an outcome (using R^2^) have relied on sequential sum of squares or stepwise regression approaches that influence the relative importance of variables and introduce bias due to researcher decisions (Grömping, U., 2007). To avoid these possibilities, we used the LMG metric from the *relaimpo* package in R (Grömping, U., 2007). LMG can account for both the direct effect an independent variable may have on the dependent variable (i.e., when the independent variable is entered into the model first), as well as its effects adjusted for other independent variables in the model (i.e., when it is entered into the model last). The LMG metric normalizes R^2^ to 100% and the contribution of each independent variable is calculated as a percentage of the R^2^ from the linear model. Using this statistical framework, we could clearly evaluate the proportion of unique variance contributed by Euclidean distance, path length, and communicability toward FC.

For each cortical parcel, we computed the R^2^ contributions for each independent variable as well as the total R^2^ contributions for each parcellation to the participant’s brain. We further mapped R^2^ for each independent variable to the participant’s functional networks.

##### 2. Convergence of S-F coupling with Margulies Unimodal-Heteromodal Gradient

Vázquez-Rodríguez et al. argued that S-F coupling follows the principal unimodal-heteromodal gradient as conceptualized by Margulies et al. (2016). Here, we tested this hypothesis explicitly.

For each participant, the principal gradient for the vertex-wise FC matrix was extracted using nonlinear dimensionality reduction via the diffusion embedding algorithm used by Margulies et al., 2016 (https://neuroanatomyandconnectivity.github.io/gradient_analysis/). These gradient maps can be observed in Fig S1. We then computed the average position along the individual-specific principal gradient for each cortical seed region.

To compare gradient position with S-F coupling, we linearly regressed the total R^2^ values obtained using the S-F regression model onto the principal gradient map obtained using the Margulies et al. approach. The residuals from this regression were used to identify regions for which S-F coupling was poorly associated with the position along the unimodal-heteromodal gradient.

## Supplementary analytic models

### Parcels derived from local gradients

To confirm that our findings were not specific to the Infomap-based algorithm as used in our primary analysis, we additionally derived parcels from a resting-state FC boundary-mapping technique (Gordon et al., 2016). This technique (based on a watershed algorithm) was then applied to each participant’s brain (Newbold et al., 2020). We generated all the previously defined variables from these parcels and re-ran our multiple linear regression model on these variables.

## Acknowledgements

This work was supported by NIH grants NS110332 (DJN), MH120989 (CJL), MH100019 (NAS), MH129493 (DMB), MH113883 (CER), MH128177 (JZ), EB031765 (JZ), DA048742 (JZ), MH120194 (JTW), NS123345 (BPK), NS098482 (BPK), MH121518 (SM), MH128696 (TX), NS124789 (SAN), MH118370 (CG), MH118362 (JRP), HD088125 (JRP), HD055741 (JRP), MH121462 (JRP), MH116961 (JRP), MH129426 (JRP), HD103525, MH120194 (JTW), MH122389 (CMS), DA047851 (CL), MH118388 (CL), MH114976 (CL), MH129616 (TOL), DA041148 (DAF), DA04112 (DAF), MH115357 (DAF), MH096773 (DAF and NUFD), MH122066 (EMG, DAF, and NUFD), MH121276 (EMG, DAF, and NUFD), MH124567 (EMG, DAF, and NUFD), NS129521 (EMG, DAF, and NUFD), and NS088590 (NUFD); by NSF grant CAREER BCS-2048066 (CG); by Center for Brain Research in Mood Disorders; by Eagles Autism Challenge; by the Dystonia Medical Research Foundation (SAN); by the National Spasmodic Dysphonia Association (EMG and SAN); by the Taylor Family Foundation (CMS, TOL); by the Intellectual and Developmental Disabilities Research Center (DJG, NUFD); by the Kiwanis Foundation (NUFD); by the Washington University Hope Center for Neurological Disorders (EMG, BPK, NUFD); and by Mallinckrodt Institute of Radiology pilot funding (DJG, EMG, NUFD).

## References

Allman, J. M., Tetreault, N. A., Hakeem, A. Y., Manaye, K. F., Semendeferi, K., Erwin, J. M., … & Hof, P. R. (2010). The von Economo neurons in frontoinsular and anterior cingulate cortex in great apes and humans. Brain Structure and Function, 214, 495–517.

Andersson, J. L., & Sotiropoulos, S. N. (2016). An integrated approach to correction for off-resonance effects and subject movement in diffusion MR imaging. Neuroimage, 125, 1063–1078.

Baum, G. L., Cui, Z., Roalf, D. R., Ciric, R., Betzel, R. F., Larsen, B., … & Satterthwaite, T. D. (2020). Development of structure–function coupling in human brain networks during youth. Proceedings of the National Academy of Sciences, 117(1), 771–778.

Braga, R. M., & Buckner, R. L. (2017). Parallel interdigitated distributed networks within the individual estimated by intrinsic functional connectivity. Neuron, 95(2), 457–471.

Brown, J. W., & Braver, T. S. (2005). Learned predictions of error likelihood in the anterior cingulate cortex. Science, 307(5712), 1118–1121.

Buckner, R. L., & Krienen, F. M. (2013). The evolution of distributed association networks in the human brain. Trends in cognitive sciences, 17(12), 648–665.

Castellanos, F. X., Margulies, D. S., Kelly, C., Uddin, L. Q., Ghaffari, M., Kirsch, A., … & Milham, M. P. (2008). Cingulate-precuneus interactions: a new locus of dysfunction in adult attention-deficit/hyperactivity disorder. Biological psychiatry, 63(3), 332–337.

Chiang, S., Stern, J. M., Engel Jr, J., & Haneef, Z. (2015). Structural–functional coupling changes in temporal lobe epilepsy. Brain research, 1616, 45–57.

Cocchi, L., Harding, I. H., Lord, A., Pantelis, C., Yucel, M., & Zalesky, A. (2014). Disruption of structure–function coupling in the schizophrenia connectome. NeuroImage: Clinical, 4, 779–787.

Damoiseaux, J. S., & Greicius, M. D. (2009). Greater than the sum of its parts: a review of studies combining structural connectivity and resting-state functional connectivity. Brain structure and function, 213, 525–533.

Fair, D.A., Miranda-Dominguez, O., Snyder, A.Z., Perrone, A., Earl, E.A., Van, A.N., Koller, J.M., Feczko, E., Tisdall, M.D., van der Kouwe, A., et al. (2020). Correction of respiratory artifacts in MRI head motion estimates. Neuroimage 208, 116400.

Fischl, B. (2012). FreeSurfer. Neuroimage 62, 774–781.

Glasser, M.F., Sotiropoulos, S.N., Wilson, J.A., Coalson, T.S., Fischl, B., Andersson, J.L., Xu, J., Jbabdi, S., Webster, M., Polimeni, J.R., et al. (2013). The minimal preprocessing pipelines for the Human Connectome Project. Neuroimage 80, 105–124. 10.1016/j.neuroimage.2013.04.127.

Goñi, J., Van Den Heuvel, M. P., Avena-Koenigsberger, A., Velez de Mendizabal, N., Betzel, R. F., Griffa, A., … & Sporns, O. (2014). Resting-brain functional connectivity predicted by analytic measures of network communication. Proceedings of the National Academy of Sciences, 111(2), 833–838.

Gordon, E. M., Laumann, T. O., Adeyemo, B., Huckins, J. F., Kelley, W. M., & Petersen, S. E. (2016). Generation and evaluation of a cortical area parcellation from resting-state correlations. Cerebral cortex, 26(1), 288–303.

Gordon, E. M., Laumann, T. O., Adeyemo, B., Gilmore, A. W., Nelson, S. M., Dosenbach, N. U., & Petersen, S. E. (2017). Individual-specific features of brain systems identified with resting state functional correlations. Neuroimage, 146, 918–939.

Gordon, E. M., Laumann, T. O., Gilmore, A. W., Newbold, D. J., Greene, D. J., Berg, J. J., … & Dosenbach, N. U. (2017). Precision functional mapping of individual human brains. Neuron, 95(4), 791–807.

Grömping, U. (2007). Relative importance for linear regression in R: the package relaimpo. Journal of statistical software, 17, 1–27.

Gusnard DA, Raichle ME. Searching for a baseline: Functional imaging and the resting human brain. Nature Reviews Neuroscience. 2001;2:685–694.

Hawellek, D. J., Hipp, J. F., Lewis, C. M., Corbetta, M., & Engel, A. K. (2011). Increased functional connectivity indicates the severity of cognitive impairment in multiple sclerosis. Proceedings of the National Academy of Sciences, 108(47), 19066–19071.

Honey, C. J., Sporns, O., Cammoun, L., Gigandet, X., Thiran, J. P., Meuli, R., & Hagmann, P. (2009). Predicting human resting-state functional connectivity from structural connectivity. Proceedings of the National Academy of Sciences, 106(6), 2035–2040.

Jackson, R. L., Bajada, C. J., Lambon Ralph, M. A., & Cloutman, L. L. (2020). The graded change in connectivity across the ventromedial prefrontal cortex reveals distinct subregions. Cerebral Cortex, 30(1), 165–180.

Jenkinson, M., Beckmann, C.F., Behrens, T.E., Woolrich, M.W., and Smith, S.M. (2012). Fsl. Neuroimage 62, 782–790.

Jeurissen, B., Descoteaux, M., Mori, S., & Leemans, A. (2019). Diffusion MRI fiber tractography of the brain. NMR in Biomedicine, 32(4), e3785.

Laumann, T. O., Gordon, E. M., Adeyemo, B., Snyder, A. Z., Joo, S. J., Chen, M. Y., … & Petersen, S. E. (2015). Functional system and areal organization of a highly sampled individual human brain. Neuron, 87(3), 657–670.

Kuceyeski, A. et al. The application of a mathematical model linking structural and functional connectomes in severe brain injury. NeuroImage: Clin. 11, 635–647 (2016).

Kuceyeski, A. F., Jamison, K. W., Owen, J. P., Raj, A. & Mukherjee, P. Longitudinal increases in structural connectome segregation and functional connectome integration are associated with better recovery after mild TBI. Hum. Brain Mapp. 40, 4441–4456 (2019).

Margulies, D. S., Ghosh, S. S., Goulas, A., Falkiewicz, M., Huntenburg, J. M., Langs, G., … & Smallwood, J. (2016). Situating the default-mode network along a principal gradient of macroscale cortical organization. Proceedings of the National Academy of Sciences, 113(44), 12574–12579.

Marcus, D.S., Harwell, J., Olsen, T., Hodge, M., Glasser, M.F., Prior, F., Jenkinson, M., Laumann, T., Curtiss, S.W., and Van Essen, D.C. (2011). Informatics and data mining tools and strategies for the human connectome project. Front. Neuroinform. 5,4.

Michel, B. F., Sambuchi, N., & Vogt, B. A. (2019). Impact of mild traumatic brain injury on cingulate functions. Handbook of Clinical Neurology, 166, 151–162.

Newbold, D. J., Laumann, T. O., Hoyt, C. R., Hampton, J. M., Montez, D. F., Raut, R. V., … & Dosenbach, N. U. (2020). Plasticity and spontaneous activity pulses in disused human brain circuits. Neuron, 107(3), 580–589.

O’Reilly, J. X., Croxson, P. L., Jbabdi, S., Sallet, J., Noonan, M. P., Mars, R. B., … & Baxter, M. G. (2013). Causal effect of disconnection lesions on interhemispheric functional connectivity in rhesus monkeys. Proceedings of the National Academy of Sciences, 110(34), 13982–13987.

Owen, J. P., Li, Y. O., Yang, F. G., Shetty, C., Bukshpun, P., Vora, S., … & Mukherjee, P. (2013). Resting-state networks and the functional connectome of the human brain in agenesis of the corpus callosum. Brain connectivity, 3(6), 547–562.

Pai, D., Muzik, O., & Hua, J. (2008, October). Quantitative analysis of diffusion tensor images across subjects using probabilistic tractography. In 2008 15th IEEE International Conference on Image Processing (pp. 1448-1451). IEEE.

Pang, J. C., Aquino, K. M., Oldehinkel, M., Robinson, P. A., Fulcher, B. D., Breakspear, M., & Fornito, A. (2023). Geometric constraints on human brain function. Nature, 1-9.

Petersen, S. E., & Sporns, O. (2015). Brain networks and cognitive architectures. Neuron, 88(1), 207–219.

Power, J.D., Barnes, K.A., Snyder, A.Z., Schlaggar, B.L., and Petersen, S.E. (2012). Spurious but systematic correlations in functional connectivity MRI networks arise from subject motion. Neuroimage 59, 2142–2154.

Power, J. D. (2017). A simple but useful way to assess fMRI scan qualities. Neuroimage, 154, 150–158.

Reveley, C., Seth, A. K., Pierpaoli, C., Silva, A. C., Yu, D., Saunders, R. C., … & Ye, F. Q. (2015). Superficial white matter fiber systems impede detection of long-range cortical connections in diffusion MR tractography. Proceedings of the National Academy of Sciences, 112(21), E2820–E2828.

Rubinov, M., & Sporns, O. (2010). Complex network measures of brain connectivity: uses and interpretations. Neuroimage, 52(3), 1059–1069.

Seeley, W. W. (2019). The salience network: a neural system for perceiving and responding to homeostatic demands. Journal of Neuroscience, 39(50), 9878–9882.

Seider, N. A., Adeyemo, B., Miller, R., Newbold, D. J., Hampton, J. M., Scheidter, K. M., … & Dosenbach, N. U. (2022). Accuracy and reliability of diffusion imaging models. NeuroImage, 254, 119138.

Smith, S.M., Jenkinson, M., Woolrich, M.W., Beckmann, C.F., Behrens, T.E.J., Johansen-Berg, H., Bannister, P.R., De Luca, M., Drobnjak, I., Flitney, D.E., et al. (2004). Advances in functional and structural MR image analysis and implementation as FSL. NeuroImage 23, S208–S219. 10.1016/j.neuroimage.2004.07.051.

Sarwar, T., Tian, Y., Yeo, B. T., Ramamohanarao, K., & Zalesky, A. (2021). Structure-function coupling in the human connectome: A machine learning approach. NeuroImage, 226, 117609.

Somers, D. C., Michalka, S. W., Tobyne, S. M., & Noyce, A. L. (2021). Individual subject approaches to mapping sensory-biased and multiple-demand regions in human frontal cortex. Current opinion in behavioral sciences, 40, 169–177.

Stephan, K. E., Friston, K. J., & Squire, L. R. (2009). Functional connectivity.

Stevens, F. L., Hurley, R. A., & Taber, K. H. (2011). Anterior cingulate cortex: unique role in cognition and emotion. The Journal of neuropsychiatry and clinical neurosciences, 23(2), 121–125.

Straathof, M., Sinke, M. R., Dijkhuizen, R. M., & Otte, W. M. (2019). A systematic review on the quantitative relationship between structural and functional network connectivity strength in mammalian brains. Journal of Cerebral Blood Flow & Metabolism, 39(2), 189–209.

Tyszka, J. M., Kennedy, D. P., Adolphs, R., & Paul, L. K. (2011). Intact bilateral resting-state networks in the absence of the corpus callosum. Journal of Neuroscience, 31(42), 15154–15162.

Van Essen, D.C., Glasser, M.F., Dierker, D.L., Harwell, J., and Coalson, T. (2012). Parcellations and Hemispheric Asymmetries of Human Cerebral Cortex Analyzed on Surface-Based Atlases. Cereb. Cortex 22, 2241–2262. 10.1093/cercor/bhr291.

Van Essen, D. C., Jbabdi, S., Sotiropoulos, S. N., Chen, C., Dikranian, K., Coalson, T., … & Glasser, M. F. (2014). Mapping connections in humans and non-human primates: aspirations and challenges for diffusion imaging. In Diffusion MRI (pp. 337–358). Academic Press.

Vázquez-Rodríguez, B., Suárez, L. E., Markello, R. D., Shafiei, G., Paquola, C., Hagmann, P., … & Misic, B. (2019). Gradients of structure–function tethering across neocortex. Proceedings of the National Academy of Sciences, 116(42), 21219–21227.

Wang, J. et al. Alterations in brain network topology and structural-functional connectome coupling relate to cognitive impairment. Front. Aging Neurosci. 10, 404 (2018).

Worrell, J. C., Rumschlag, J., Betzel, R. F., Sporns, O., & Mišić, B. (2017). Optimized connectome architecture for sensory-motor integration. Network Neuroscience, 1(4), 415–430.

Zhang, Y., Brady, M., & Smith, S. (2001). Segmentation of brain MR images through a hidden Markov random field model and the expectation-maximization algorithm. IEEE transactions on medical imaging, 20(1), 45–57.

