## Supplementary Figures for "Structure-Function Coupling in Highly Sampled Individual Brains"

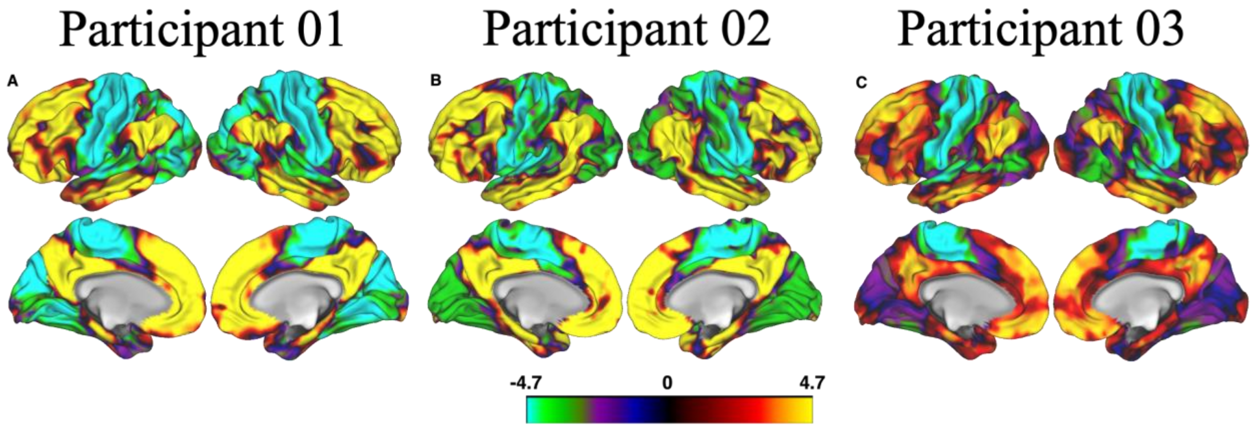


**Figure S1: First principal gradient values derived from the Margulies et al. 2016 diffusion-embedding algorithm**. ***Panels A-C*** *correspond to participant 01(A),02(B), and 03(C). Principal gradient values were thresholded between -4.7(unimodal areas) to 4.7(multimodal areas) for easy comparison across participants.*

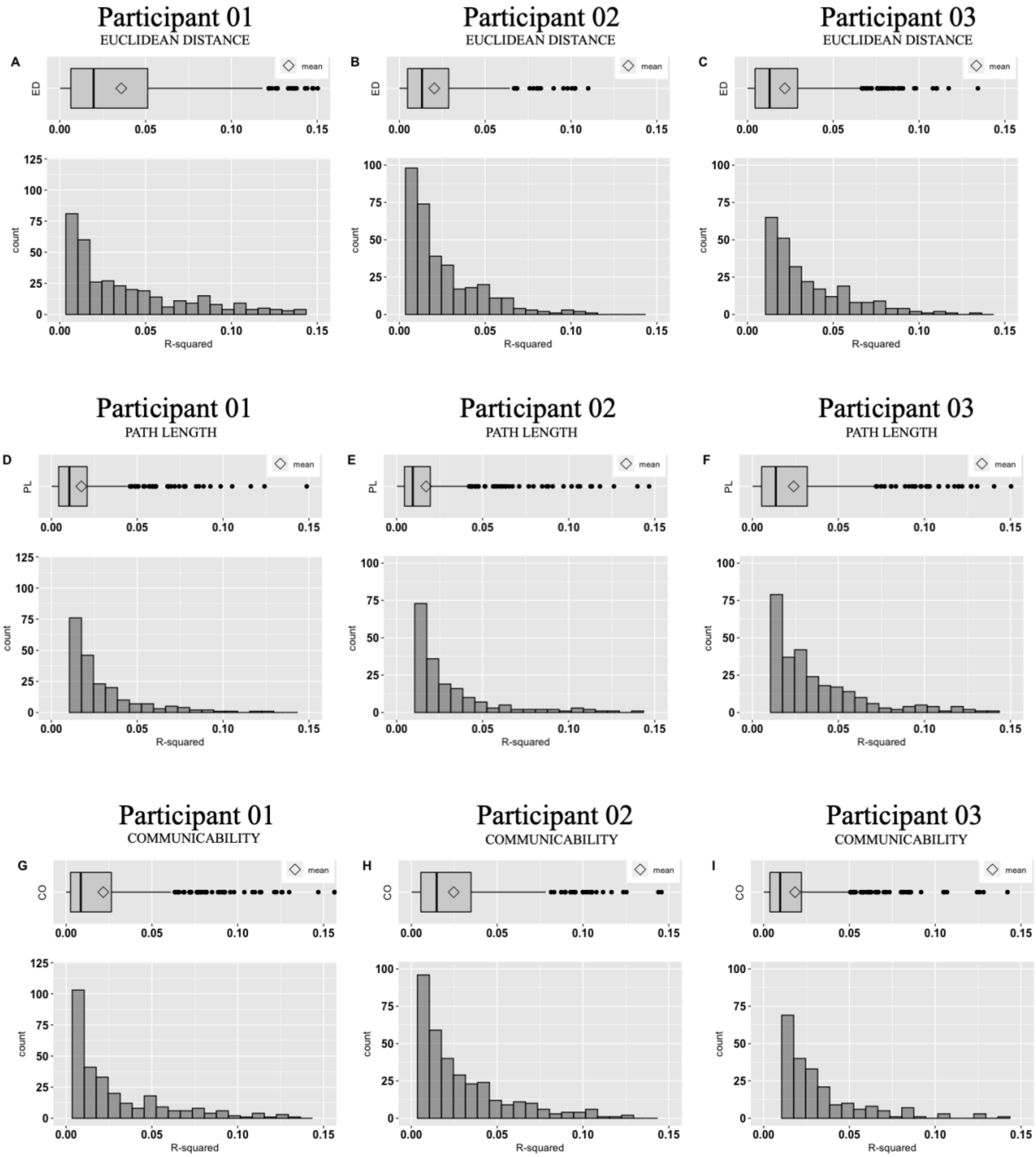


**Figure S2: Distribution of variance contributions depicted by boxplots and histogram plots for each participant's Infomap algorithm-derived brain organization**. ***Panels A-C****: Boxplot and histogram plot for variance contribution by* ***Euclidean distance*** *toward functional connectivity (FC) for participant 01(A),02(B), and 03(C) respectively.* ***Panels D-F****: Boxplot and histogram plot for variance contribution by (weighted)* ***path length*** *toward FC for participant 01(D),02(E), and 03(F) respectively.* ***Panels G-I****: Boxplot and histogram plot for variance contribution by (weighted)* ***communicability*** *toward FC for participant 01(G),02(H), and 03(I) respectively. Across all panels,* $R^2$ *values were scaled to 0.15 maximum comparability across participants.*

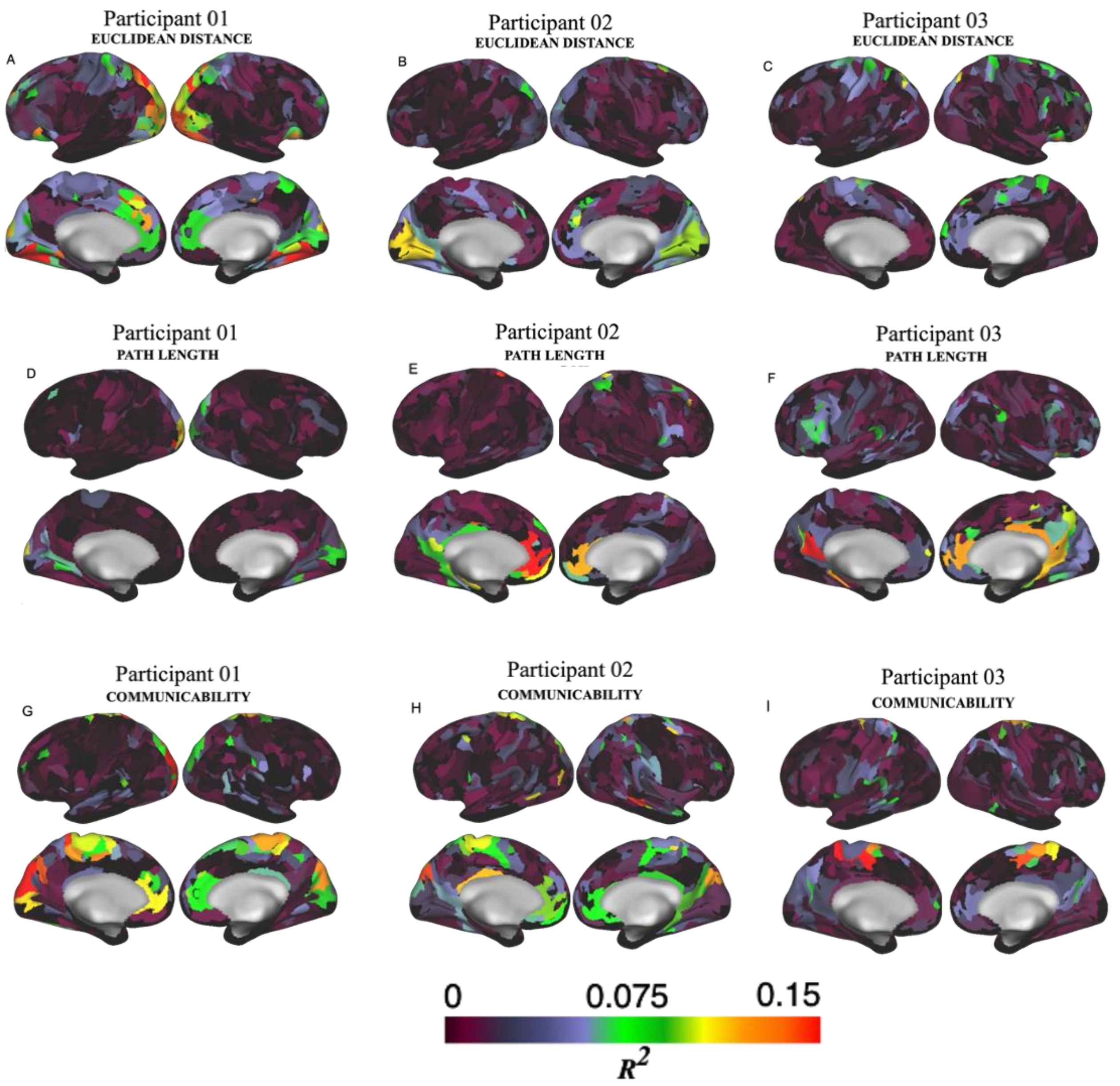


**Figure S3: Variance contributions are mapped onto each participant's specific brain organization, derived using the Infomap community-driven algorithm**. ***Panels A-C:*** *Variance contribution by* ***Euclidean distance*** *toward FC for participant 01(A),02(B), and 03(C) respectively.* ***Panels D-F****: Variance contribution by (weighted)* ***path length*** *toward FC for participant 01(D),02(E), and 03(F) respectively.* ***Panels G-I****: Variance contribution by (weighted)* ***communicability*** *for participant 01(G),02(H), and 03(I) respectively. $R^2$ values were thresholded to 0.15 for easy comparison across participants. Black color indicates $R^2$ values = 0.*

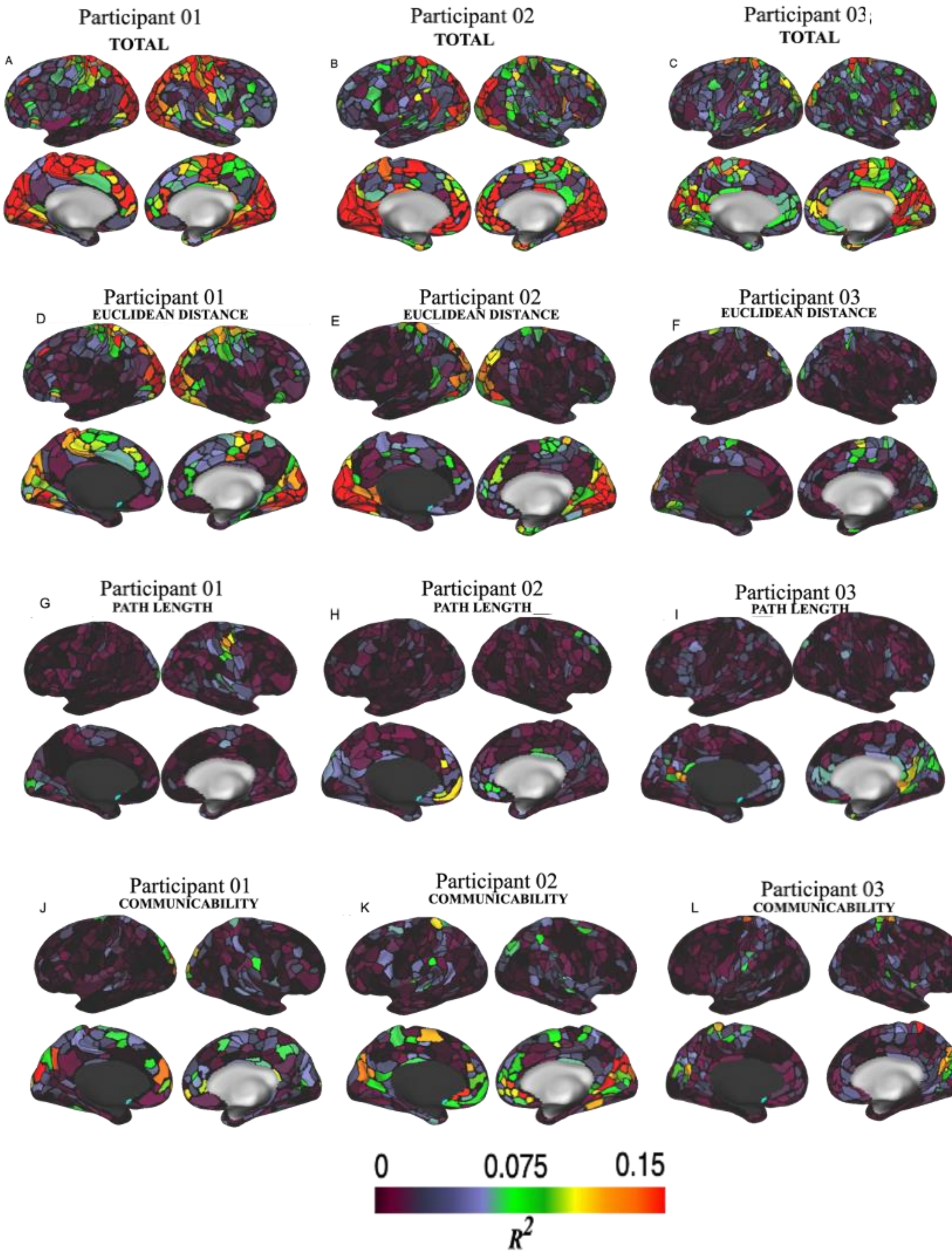


**Figure S4: Variance contributions are mapped onto each participant's specific brain organization, derived using the watershed algorithm**. ***Panels A-C***: ***Total*** *variance contribution by structural metrics toward FC for participant 01(A),02(B), and 03(C).* ***Panels D-F***: *Variance contribution by* ***Euclidean distance*** *toward FC for participant 01(D),02(E), and 03(F).* ***Panels G-I***: *Variance contribution by (weighted)* ***path length*** *toward FC for participant 01(G),02(H), and 03(I) respectively).* ***Panels J-L***: *Variance contribution by (weighted)* ***communicability*** *for participant 01(J),02(K), and 03(L) respectively).*

*$R^2$ values were thresholded to 0.15 for easy comparison across participants. Black color indicates $R^2$ values = 0.*

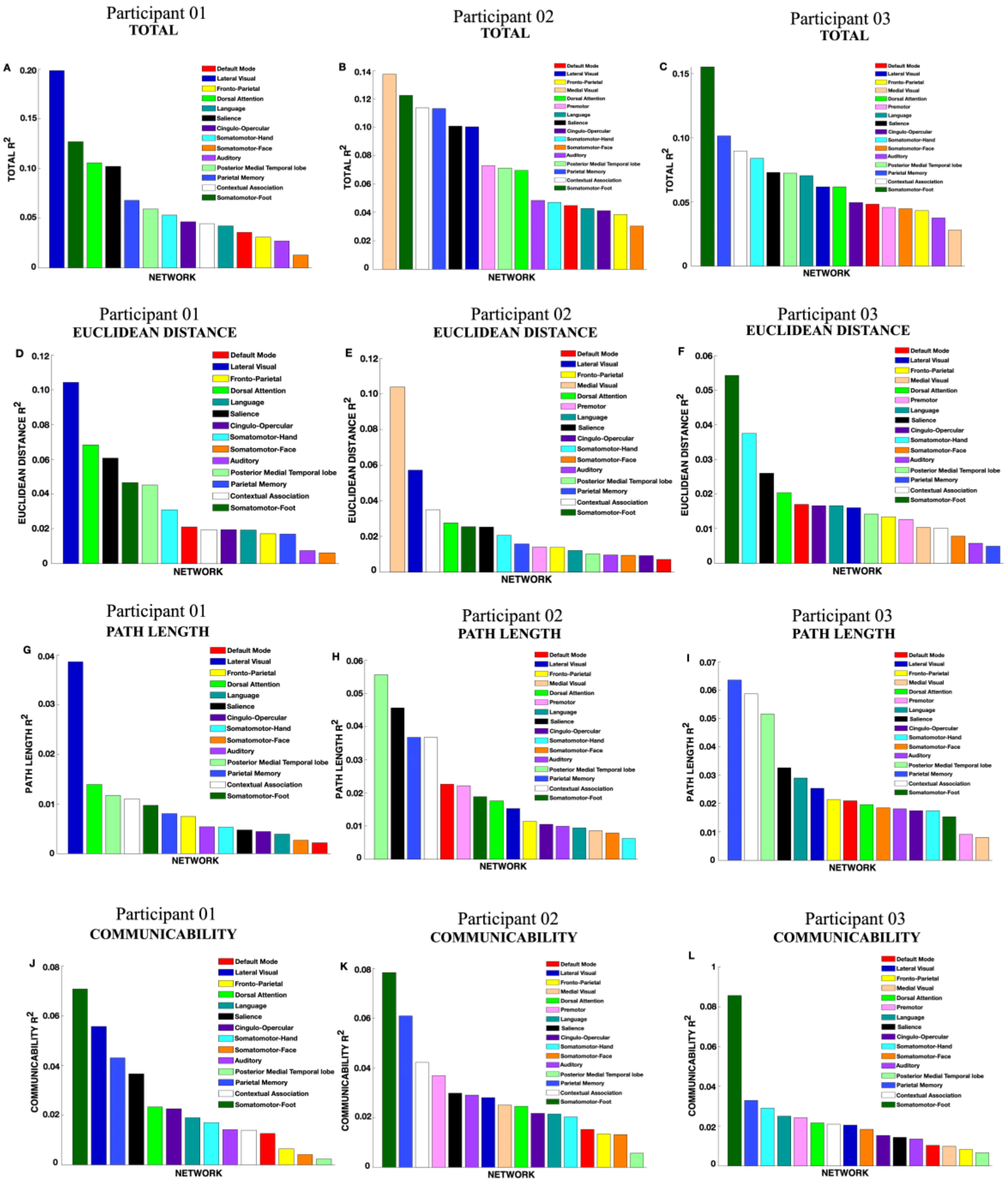


**Figure S5: Variance contributions toward parcels affiliated with 14 resting state FC networks for participant 01, and 16 resting state FC networks for participants 02 and 03**. ***Panels A-C***: ***Total*** *variance contribution by structural metrics toward parcels affiliated with resting state FC networks for participant 01(A),02(B), and 03(C).* ***Panels D-F***: *Variance contribution by* ***Euclidean distance*** *toward parcels affiliated with resting state FC networks for participant 01(D),02(E), and 03(F).* ***Panels G-I***: *Variance contribution by (weighted)* ***path length*** *toward parcels affiliated with resting state FC networks for participant 01(G),02(H), and 03(I).* ***Panels J-L***: *Variance contribution by (weighted)* ***communicability*** *toward parcels affiliated with resting state FC networks for participant 01(J),02(K), and 03(L).*
